# Transcranial direct current stimulation of the supplementary motor area modulates behavioral and peripheral electrophysiological measures of motor preparation

**DOI:** 10.1101/2025.10.31.685710

**Authors:** Ashutosh Kumar, Ayusha Desai, Kalpajyoti Hazarika, Aditya Murthy

**Author notes:** Corresponding author: Aditya Murthy, PhD, Movement Control Lab, Centre for Neuroscience, Indian Institute of Science CV Raman Road, Bengaluru 560012, India.

## Abstract

Converging evidence has shown that the supplementary motor area (SMA) is involved in the preparation of movements. Such preparatory activity has traditionally been thought to lead to shorter reaction times by allowing the motor network to settle into states favourable to motor execution without directly engaging the peripheral musculature, thus limiting preparatory states to central neural processes. We revisited this issue by recording motor unit activity using high-density surface EMG grid electrodes from a proximal muscle, the anterior deltoid, during a delayed reach task. We observed that anodal stimulation of the SMA enhances motor preparation by shortening reaction times while simultaneously increasing motor unit activity. Additionally, we found that SMA stimulation produced a subsequent increase in movement velocities due to the activation of larger-amplitude motor units. Taken together, these findings suggest that motor preparatory states involve an active interaction between central and peripheral processes. Direct descending SMA outputs to the spinal cord provide a link between cortical preparation and muscle readiness, resulting in increased motor vigor.

## Introduction

The execution of skilled voluntary movements relies on motor preparation, which involves organising the motor network into states that favour subsequent movement execution (Shenoy et al., 2013; Churchland et al., 2006,2010,2012; Kaufman et al., 2014; Elsayed et al., 2016). Studies have also shown that the Supplementary Motor Area (SMA) is a key node in the network that orchestrates this preparatory process (Tanji, 1994; Picard & Strick, 1996; Nachev et al., 2008), with convergent human evidence from MRI and ERP/EEG demonstrating SMA/pre-SMA engagement during readiness and initiation (Tanji,1984; Cunnington et al., 2002; Jenkins et al., 2000; Shibasaki & Hallett, 2006; Lau et al., 2006). For example, electrophysiological studies in non-human primates show that neurons in medial premotor cortex (SMA/pre-SMA) exhibit robust delay-period activity that ramps during preparation and anticipates movement onset, and ceases firing just prior to execution, distinguishing preparatory and execution phases (Tanji & Shima, 1994; Tanji, 1994; Crammond & Kalaska, 2000; Tanji & Kurata, 1982,1985; Shima & Tanji, 2000; Tanji, 2001). This temporal signature positions the SMA as a critical locus for establishing a preparatory motor set—an active neurophysiological state that optimizes initial cortical conditions, thereby preconfiguring the downstream motor network for the rapid execution of voluntary movement (Carlson et al., 2015; Churchland et al., 2012; Nachev et al., 2008; Picard & Strick, 1996; Shenoy et al., 2013; Tanji, 1994). Human EEG studies further confirm this role, identifying slow-rising movement-related potentials over medial frontal cortex/SMA that begin well before movement execution, including the classic Bereitschaftspotential and its early and late components (Shibasaki & Hallett, 2006; Deecke & Kornhuber, 1978; Cunnington et al., 2002; Colebatch, 2007).

The behavioral relevance of preparatory activity is evident in reaction time paradigms, where introducing a delay period between target appearance and go cue consistently results in faster response times - the "hold time effect" (Churchland et al., 2006,2010; Shenoy et al., 2013; Lara et al., 2018). Systematic manipulations of the delay period duration, combined with transcranial magnetic stimulation (TMS), have been used to map how inhibitory processes dynamically fluctuate during motor preparation (Lebon et al.,2015). Studies applying anodal tDCS over the SMA have demonstrated polarity-dependent effects on reaction time performance, consistent with modulating preparatory activation levels toward an action threshold (Carlsen et al., 2015; Sadler et al., 2022; Sato et al., 2024).

Despite the presence of the hold time effect, traditional EMG methods have largely failed to detect muscle activity during preparatory periods (Tanji & Evarts, 1976; Kaufmann et al., 2014), leading to the notion that preparatory states are confined to central neural processes. However, early work by Mellah et al. (1990) showed changes in motor unit excitability during movement preparation in non-human primates, suggesting that preparatory processes might reach the peripheral musculature. In addition, recent work by Rungta and Murthy (2023) used high-density surface EMG to show that early motor preparation can be detected peripherally through context-specific recruitment of small-amplitude motor units during the delay period in reaching tasks in humans. These "ramping" motor units displayed spatially specific activity patterns that correlated with reaction times, providing direct evidence of preparatory neural drive extending to peripheral neuromuscular readiness.

The presence of peripheral preparatory signatures raises a crucial question: do cortical neuromodulation effects, such as those induced by SMA tDCS, translate to measurable changes in the early recruitment patterns of these preparatory motor units? The present study addresses this question by investigating whether anodal tDCS applied to the SMA modulates the early preparatory activity of low-threshold motor units in the anterior deltoid muscle during a randomized delayed reach task. By combining targeted cortical stimulation with high-density EMG recordings, we aimed to establish a direct link between cortical neuromodulation and peripheral manifestations of motor preparation.

## Materials and Methods

### Participants

Thirty-seven healthy, right-handed volunteers (*N* = 37; age range: 19–26 years) participated in the experiment and were included in the final analysis. All participants reported normal or corrected-to-normal vision, no history of neurological or psychiatric disorders, and no concurrent use of medication affecting the central nervous system. Written informed consent was obtained from all participants prior to the experiment, in accordance with the protocol approved by the Institutional Human Ethics Committee of the Indian Institute of Science, Bengaluru. Participants received monetary compensation for their time.

The final analyzed cohort was divided into two independent experimental groups: a Test group (*n* = 23) and a Sham group (*n* = 14). Participants in the Test group underwent active anodal tDCS, with the anode positioned over the supplementary motor area (SMA). Participants in the Sham group underwent an identical experimental setup and electrode montage; however, they received sham stimulation without sustained active current delivery. Across all initial recruitments, one individual was systematically excluded from the final pooled sample due to failure to exhibit the hold-time effect. This subject also demonstrated poor task performance (e.g., executing correct responses in fewer than 50 out of 100 trials across conditions).

### Experiment Setup

Stimulus presentation and data collection were conducted using NIMH MonkeyLogic (Hwang et al., 2019) in MATLAB (MathWorks Inc.). Visual stimuli appeared on a 21.5-inch XP Pen Artist Display 22R Pro tablet (Hanvon Ugee Group, Shenzhen, China) operating at a resolution of 1920 × 1080 pixels at 60 Hz. This device also functioned as the primary response interface, featuring a pressure-sensitive, battery-free stylus (8192 pressure levels, 5080 LPI resolution, tilt sensitivity). For a subset of participants, the visual task was mirrored on a separate external monitor with identical resolution and refresh rate specifications. In these cases, the subjects still executed all physical reaching movements exclusively on the XP Pen tablet. This dual-display configuration introduced no technical latency or hardware discrepancy to the data collection pipeline. Regardless of the specific visual setup, participants moved the stylus directly across the tablet surface, and the continuous positional and kinematic data were recorded by the MonkeyLogic software at a strict 1 kHz sampling rate for offline analysis.

### Task Design

Participants performed a randomized reach paradigm comprising Delayed Reaction Time (DRT) trials and Simple Reaction Time (SRT) trials. In both trial types, participants began by positioning the stylus at a central fixation point (FP) displayed on an XP-Pen tablet. In the Delayed RT trials, following a 500 ms fixation period, a green peripheral target (a square) appeared at one of two possible locations, while the fixation point remained illuminated for an additional 1000 ms (Hold Time). During this hold period, participants were required to withhold movement despite the target being visible. Participants were instructed to initiate their reaching movement towards the target only after the FP disappeared, which served as the Go cue (Figure 1, left). This delay period allowed us to dissociate target selection and motor preparation processes from movement execution. For Simple RT trials (Figure 1, right), the target onset coincided with the immediate disappearance of the FP, prompting participants to initiate movement as quickly as possible without any imposed delay. This condition served as a baseline or control for measuring reaction times without an intervening hold period. Both trial types were randomly intermixed in equal proportions throughout each session to prevent anticipatory motor planning. Each session comprised 100 experimental trials (50 trials to each of two predefined target positions: ±4.5° horizontal, ±11° vertical from the central FP). Successful target acquisition within a predetermined time window was signalled with an auditory feedback tone (1000 Hz, 100 ms duration). To ensure familiarity with the task requirements and reduce learning effects during data collection, participants completed 20–25 practice trials before beginning the main experimental session.

**Figure 1.**
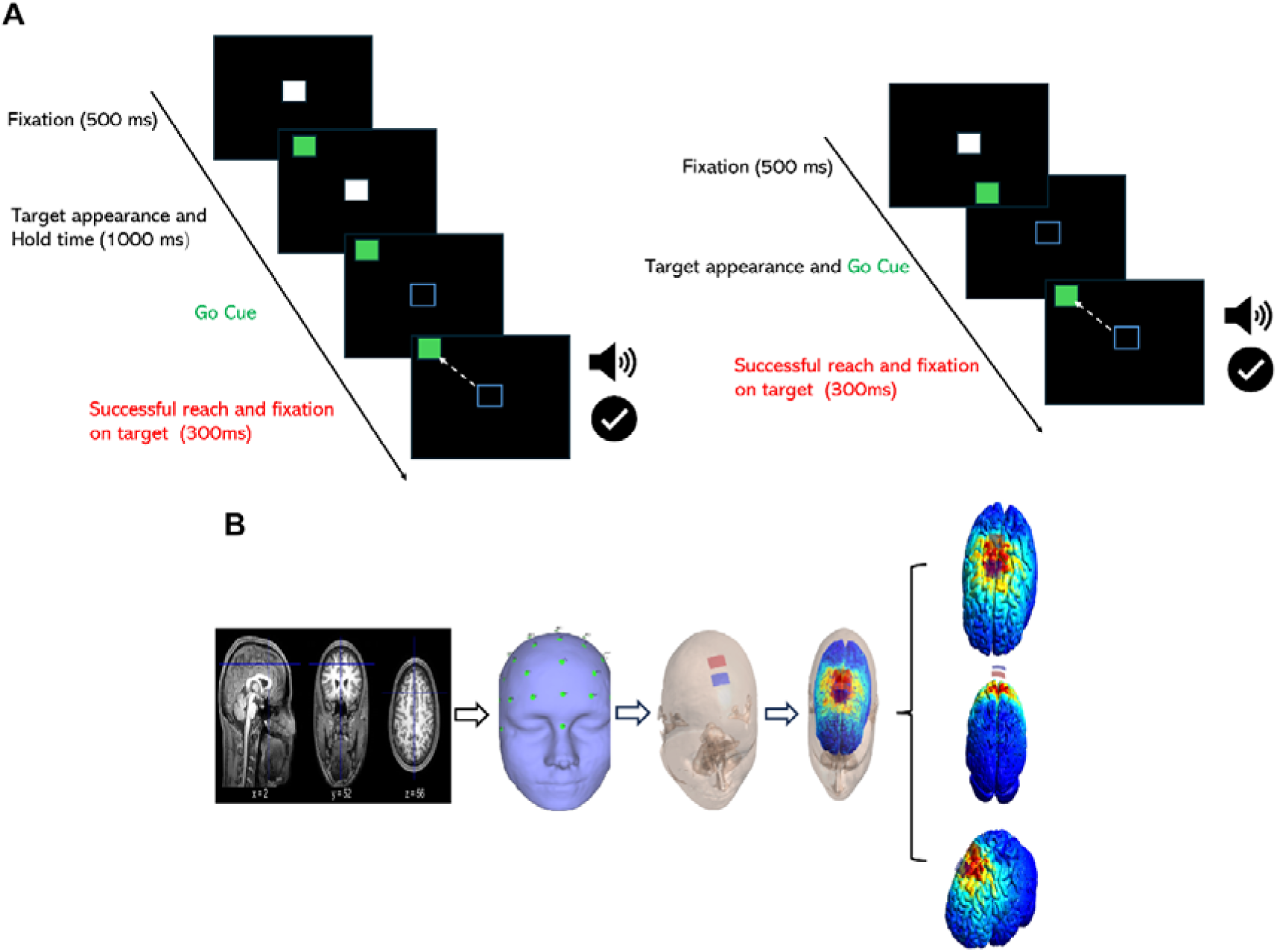
Schematic diagram of task paradigm. (A) Task design: Delayed Reaction Time trials (left) with fixation, target appearance, and delayed go cue; Simple Reaction Time trials (right) with simultaneous target onset and go cue. (B) Head modelling pipeline: MRI acquisition, 3D reconstruction and tissue segmentation, electrode placement and electric field simulation results showing current density distributions.

### tDCS Protocol

#### Device and Electrode Configuration

Transcranial direct current stimulation was delivered using a 1×1 low intensity stimulator (Model 2004, Soterix Medical Inc). Custom carbon rubber electrodes (2.2 ×3 cm; surface area 6.6 cm²) coated with electrode conductive paste were used to ensure durability, stable conductivity, and improved focality compared to larger sponge electrodes, thereby reducing current spread to adjacent brain regions.

#### Stimulation location and parameters

For one participant, SMA was localised anatomically from individual T1-weighted MRI scans (1 mm isotropic voxel resolution). The estimated location of the SMA for this participant was estimated as being 1.8 cm anterior to Cz, which was consistent with previous studies (Carlsen et al., 2015; Sadler et al., 2022). Therefore, in the remaining participants, the SMA location was determined using the international 10–20 EEG system, defined as 1.8 cm anterior to Cz. In the anodal condition, the anode was placed over SMA and the cathode over FCz. Stimulation was delivered at 2.0 mA for 10 minutes with 30 s ramp-up and ramp-down. In the sham condition, electrodes were positioned identically, but stimulation was limited to a brief 30 s ramp-up and ramp-down at the start to mimic sensation without the current delivery. After a pre-stimulation behavioural session, participants received stimulation according to protocol, rested for 10 minutes, and then performed the post-stimulation session.

#### Electric Field and Current Distribution Simulation

With high-resolution T1-weighted MRI scans, a finite element head model was generated using the SimNIBS 4.1 “CHARM” pipeline (Puonti et al., 2020) (Figure 1B). The pipeline segmented scalp, skull, cerebrospinal fluid, gray matter, and white matter, and incorporated diffusion data when available to model tissue anisotropy. Electrode size, location, polarity, and stimulation parameters matched those used in the experiment (2 mA; 2.2× 3 cm electrode; SMA target 1.8 cm anterior to Cz; return at FCz). Head meshes segmented scalp, skull, CSF, gray, and white matter; electrode size, location, and current matched the experimental setup. Simulations produced spatial maps of induced electric fields (V/m) and current density (A/m²), allowing assessment of focality and peak field magnitude within SMA. (see Figure 1B).

### Data Acquisition and Analysis

#### Behaviour Data

Continuous two-dimensional stylus trajectories (x, y) were sampled at 1 kHz using MonkeyLogic, a MATLAB-based real-time behavioral control and data acquisition suite (Hwang et al., 2019). Offline trajectory smoothing was achieved using a 5-point moving average filter applied with zero-phase distortion(filfilt) to eliminate directional temporal shifts. Tangential velocity and acceleration profiles were subsequently derived via first- and second-order discrete numerical differentiation of the filtered spatial coordinates.

Reaction time (RT) was defined as the temporal interval from the presentation of the Go cue to overt movement onset. Movement onset was taken as the timestamp at which the tangential reaching velocity first exceeded 5 cm/s. To isolate and quantify the behavioral cost of motor preparation, we calculated a Hold Time Effect for each participant. This metric was defined as the within-subject difference in reaction time between delayed (prepared) and simple (unprepared) reach trials:

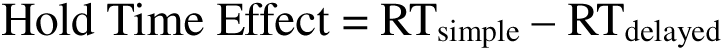

Execution kinematics, specifically peak reaching velocity and maximum acceleration, were extracted directly from the differentiated temporal profiles to serve as metrics of movement kinematics. To guarantee the integrity of the behavioral dataset, rigorous quality control criteria were enforced prior to statistical evaluation. Trials exhibiting anticipatory responses (RT < 100 ms), missed targets, or gross trajectory deviations were systematically excluded from all downstream analyses.

### EMG Signal recording, Pre-processing, and Motor Unit Analysis

All signal processing, statistical analyses, and data visualization were executed using custom-developed algorithms in MATLAB (MathWorks, Inc.). High-density surface electromyography (HD-sEMG) signals were acquired from the anterior deltoid at a sampling frequency of 2000 Hz using a Muovi system (OT Bioelectronica, Italy). The recording array consisted of 32 electrodes spaced 5 mm apart, configured in a 8×4 grid. To enhance the signal-to-noise ratio (SNR) and suppress common-mode interference, the monopolar recordings were mathematically differentiated into 28 bipolar channels (7 rows × 4 columns) along the direction of the muscle fibers.

Raw bipolar signals were pre-processed, with a fourth-order Butterworth bandpass filter (10– 500 Hz) to eliminate low-frequency movement artifacts and high-frequency noise, followed by standard spectrum estimation techniques to remove line noise. To isolate the most robust representation of global muscle activation, the single bipolar channel exhibiting the highest SNR—defined by the maximal decibel increase from the pre-movement baseline (1000 ms before go cue) to the active execution phase (1000 ms after go cue)—was selected for further analysis. Continuous gross EMG amplitude envelopes were extracted by computing the root-mean-square (RMS) using a 25 ms sliding temporal window to capture transient neuromuscular bursts. To systematically account for inter-subject impedance variability and enable unbiased cohort comparisons, individual RMS envelopes were normalized to each subject’s maximum mean task-evoked amplitude (In-RF peak execution). This yielded standardized execution trajectories scaled from 0 to 1, preserving relative activation dynamics while mitigating absolute amplitude artifacts. Trials containing missing values or artifacts were systematically excluded.

#### Motor Unit Decomposition and Firing Rate Extraction

Putative single-motor unit action potentials were discriminated using an adaptive peak detection algorithm. Baseline electrophysiological activity was computed from a 500 ms quiescent window during the hold time window of the task. To account for the size-dependent amplitude scaling of the motor pool, detection thresholds were set at 3, 5, and 7 standard deviations above the baseline mean for small, medium, and large putative motor units, respectively (Rungta and Murthy, 2023). Hereafter, these amplitude-discriminated putative units are referred to simply as motor units throughout the remainder of this study.. Robust spike identification was enforced via strict temporal constraints, including a minimum inter-spike interval of 40 ms and peak-width boundaries of 10 to 40 ms.

To estimate continuous instantaneous firing rates, discrete spike trains were converted into peri-event time histograms (PSTHs) utilizing 10 ms temporal bins. The histograms were temporally aligned to both the Go cue (for preparation) and movement onset (for execution). To prevent temporal phase distortion during smoothing, the binned spike counts were convolved with a Gaussian kernel (sigma = 20 ms) utilizing zero-phase forward and reverse digital filtering (filtfilt). Firing rates were subsequently averaged across valid trials for each condition. To allow for direct comparisons of recruitment thresholds across the motor pool size hierarchy, continuous firing rates were normalized to each specific unit’s absolute peak execution firing rate.

#### Population-Level Analysis

For test cohort-level (n = 23), firing rates and RMS amplitudes were averaged over two pre- defined 500 ms temporal epochs: a preparation window (−0.50 to 0.00 s relative to the Go cue) and an execution window (0.00 to +0.50 s relative to movement onset). This yielded a single, paired mean value per participant, per epoch, per condition (e.g., In-RF vs. Out-RF; Pre- vs. Post-stimulation). To investigate the relationship between preparatory and execution states, a Modulation Index (MI) was calculated for each participant

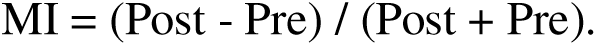

where *Post* and *Pre* represent the normalized mean firing rates extracted from the respective epoch. This formulation yields a standardized metric strictly bounded between -1 and 1, where positive values indicate a stimulation-induced facilitation of motor unit discharge, and negative values indicate suppression. By converting absolute firing rates into this relative index, we were able to directly correlate the degree of preparatory modulation with execution-phase dynamics across the population.

### Statistical Analysis

Statistical analyses were performed in MATLAB (MathWorks, Inc.) using the Statistics and Machine Learning Toolbox. Data normality was assessed with the Shapiro-Wilk test to determine the appropriate use of parametric or non-parametric tests.

Behavioral metrics, including reaction time (RT) and peak movement velocity, were analyzed using paired within-subject comparisons (*n* = 23). Based on previous literature, we used one-tailed tests to evaluate directional hypotheses, expecting active tDCS to decrease RT and increase peak reaching velocity. We used paired-sample *t*-tests for normally distributed data and Wilcoxon signed-rank tests when the paired differences deviated from normality.

For single-subject EMG analysis, firing rates were extracted trial-by-trial. Since the number of valid trials naturally varied between pre- and post-stimulation sessions, these trial distributions were treated as independent samples. We evaluated differences in single-subject firing rates using either the independent two-sample *t*-test or the Wilcoxon rank-sum test (Mann-Whitney *U* test), depending on the normality results.

For population-level comparisons, time-series data were averaged into discrete values. Firing rates and gross EMG RMS amplitudes were averaged over two 500 ms epochs: a preparation window (−0.50 to 0.00 s relative to the Go cue) and an execution window (0.00 to +0.50 s relative to movement onset). This provided a single paired mean value per participant, epoch, and condition. Population-level differences were then evaluated using paired-sample *t*-tests or Wilcoxon signed-rank tests. To examine linear relationships between preparatory and execution states, we computed a Modulation Index (MI) and Pearson correlations was used to test the linearity between two phases. Statistical significance was set at α = 0.05. Effect sizes are reported alongside standard inferential statistics: Cohen’s *d* for parametric comparisons and the rank-biserial correlation coefficient (r_rb_) for non-parametric tests. Exact *p*-values, relevant test statistics, effect sizes, and sample sizes are detailed throughout the text and figure legends.

## Results

### tDCS over SMA accelerates motor initiation and enhances execution kinematics

To determine whether the neurophysiological changes induced by transcranial direct current stimulation (tDCS) translate to behavioral motor performance, we evaluated the hold time effect (the reaction time difference between delayed and simple trials) and maximum reaching velocity across the cohort (Test, n = 23; Control, n = 14). In the test group, tDCS accelerated motor initiation. The hold time effect significantly decreased post-stimulation (Pre: Median = 170.00, IQR = 165.75 vs. Post: Median = 119.00, IQR = 129.62 ms; Wilcoxon signed-rank test, W = 37.00, one-tailed p = 0.003, rank biserial correlation = −0.57). Importantly, the hold time effect was absent in the sham-control group (Pre: Median = 102.00, IQR = 136.00 vs. Post: Median = 80.75, IQR = 127.50 ms; Wilcoxon signed-rank test, W = 29.00, one-tailed p = 0.073), confirming the specificity of the neuromodulatory intervention of tDCS (Figure 2A). Having established that active tDCS accelerates motor initiation, we next investigated whether this intervention causally modulates the kinematics of the subsequent physical movement. To quantify execution kinematics, we evaluated the mean peak reaching velocity (cm/s) across both the cohorts. Because paired differences in mean peak velocity significantly deviated from normality (Shapiro-Wilk test, *p* = 0.010 for both cohorts), we used exact, non-parametric statistics for all within-group hypothesis testing. Consistent with the observed facilitation in motor onset, the test cohort (*n* = 23) demonstrated a significant enhancement in execution kinematics following active neuromodulation. Peak execution velocities increased from a pre-intervention median of 41.55 cm/s (IQR = 11.05) to a post-intervention median of 42.22 cm/s (IQR = 12.55). This facilitation yielded a median intra-subject kinematic gain of 2.49 cm/s (IQR = 3.63 cm/s), confirming a significant, directional increase in motor output (Wilcoxon signed-rank test, one-tailed; *z* = 2.966, *p* = 0.0015) (Figure 2B). This behavioural modulation was driven by a large effect size (Rank Biserial Correlation = 0.618) and achieved robust statistical power (0.840). In contrast, the sham Control cohort (*n* = 14) demonstrated no such kinematic enhancement. Peak execution velocities in the control group failed to increase following the mock intervention (Pre: Median = 48.37, IQR = 12.26 cm/s vs. Post: Median = 42.78, IQR = 14.55 cm/s), resulting in a negligible median paired difference of just 0.38 cm/s (IQR = 6.93 cm/s). Statistical evaluation confirmed the complete absence of a stimulation-induced increase in kinematics (Wilcoxon signed-rank test, one-tailed; *p* = 0.729), characterized by a small, negative effect size (Rank Biserial Correlation = -0.181).

**Figure 2.**
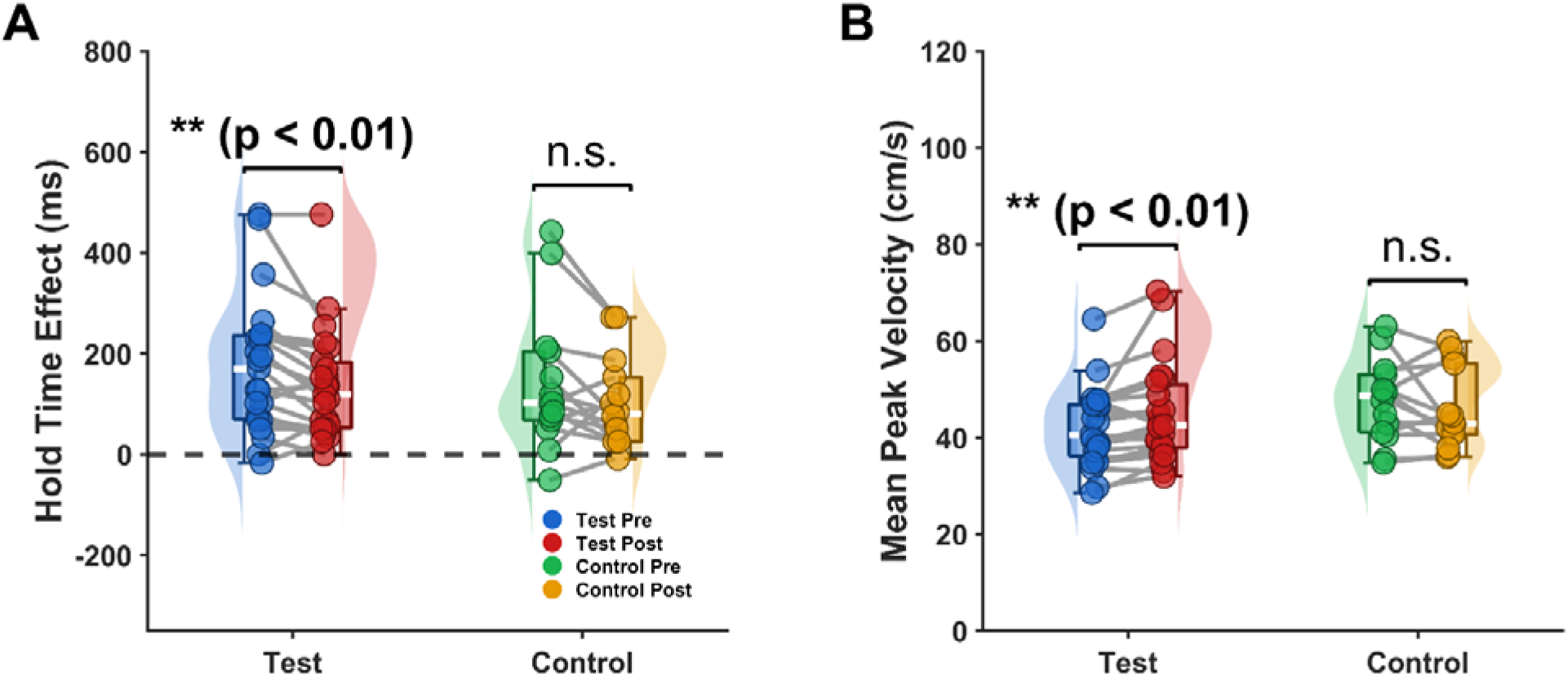
Active neuromodulation accelerates motor initiation and enhances execution kinematics. (A) Modulation of motor initiation. Raincloud plots illustrating the hold time effect (quantified as the reaction time difference between delayed and simple reach trials) before (Pre) and after (Post) the intervention. The active tDCS Test group (blue/red; 23 ) exhibited a significant reduction in the hold time effect post-stimulation, indicating accelerated motor initiation (Wilcoxon signed-rank test, *p* < 0.01). In contrast, the sham Control group (green/yellow; 14 ) showed no significant behavioral modulation (n.s.). The dashed horizontal line indicates a net zero hold time effect. (B) Enhancement of execution kinematics. Population-level paired comparisons of mean peak reaching velocity (cm/s). Consistent with the initiation metric, the test cohort demonstrated a statistically significant elevation in execution vigor following active stimulation (*p* < 0.01). Kinematic parameters in the sham/control cohort remained statistically invariant (n.s.). For both panels, flanked half-violins denote the probability density of the behavioral distributions. Internal box plots display the sample median (thick white horizontal line) and interquartile range (IQR, colored box). Connected scatter points represent the paired trajectory of individual participants across the experimental phases.

### High-density motor unit decomposition unmasks spatially tuned preparatory drive and robust execution dynamics

To determine whether motor preparatory commands activate the peripheral musculature prior to active movement, we examined anterior deltoid activation across all test participants (*n* = 23) during the task. We first analyzed the global muscle output using conventional root-mean-square (RMS) EMG aligned to movement onset. For movements directed into the response field (In-RF; target at −4.5°, +11°), global RMS EMG exhibited robust, phasic activation tightly coupled to movement execution. Interestingly, this signal remained flat throughout the preceding delay period. This lack of observable delay activity aligns with the classical consensus that standard surface EMG cannot detect preparatory neural dynamics prior to movement onset (**Figure 3A**).

**Figure 3.**
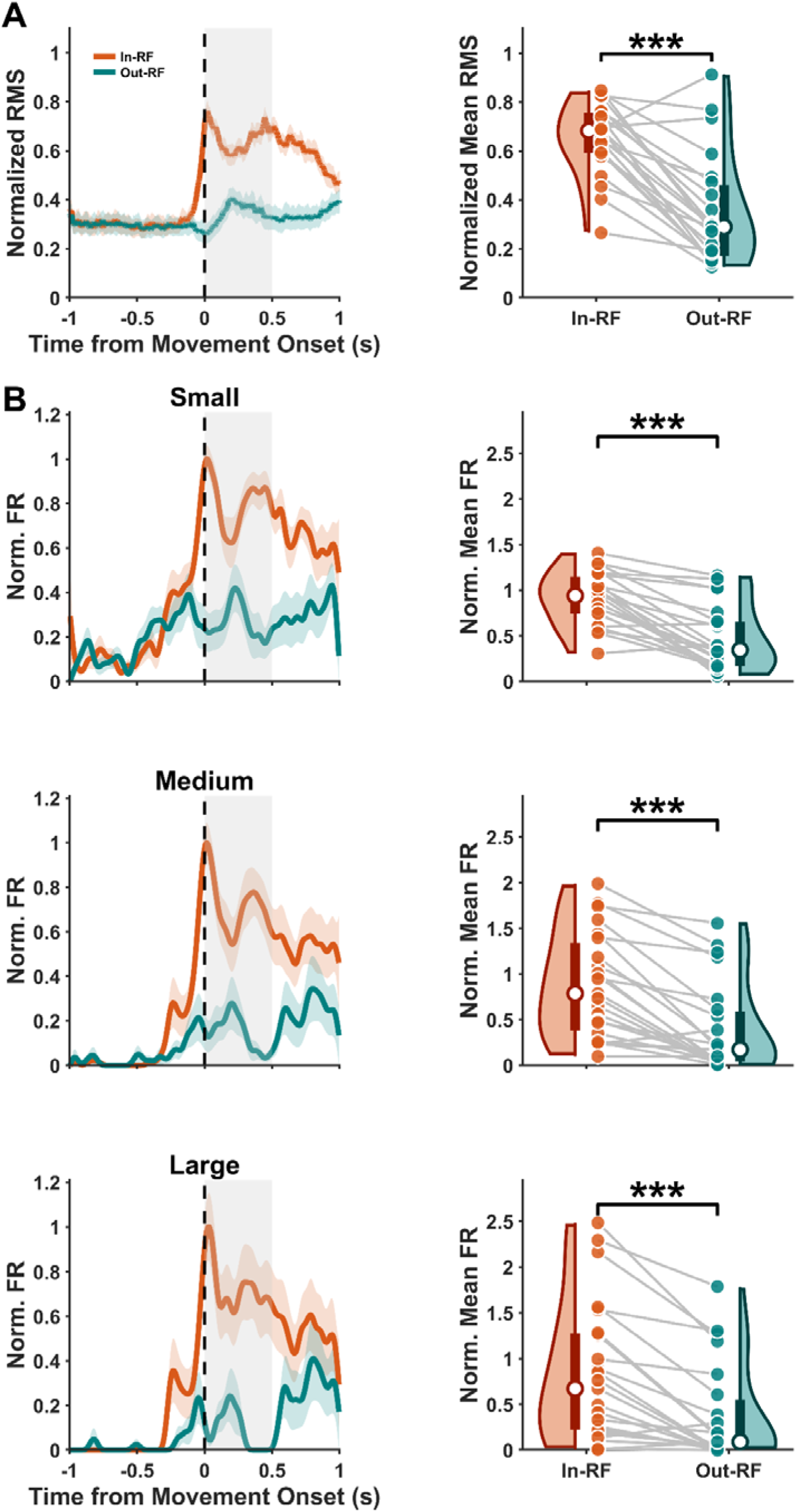
Spatial tuning of gross muscle activation and hierarchical motor unit recruitment during active movement execution. **(A)** *Left:* Population-averaged continuous trajectories (*N* = 23) of normalized RMS EMG from the anterior deltoid, aligned to movement onset (dashed vertical line). Movements directed into the receptive field (In-RF; orange) exhibit a robust, phasic amplification during execution compared to non-preferred trajectories (Out-RF; teal). Solid lines indicate the population mean; shaded temporal envelopes denote the standard error of the mean (SEM). The gray shaded epoch (0.00 to 0.50 s) delineates the execution window utilized for statistical quantification. *Right:* Raincloud plots demonstrating population-level paired differences in normalized mean RMS during the execution epoch. Half-violins represent the probability density of the data, internal box plots indicate the median and interquartile range (IQR, with the white dot representing the median), and connected scatter points represent paired individual participant means (Wilcoxon signed-rank test, *** *p* < 0.001). **(B)** Directional tuning dynamics mapped across the motor unit size hierarchy. *Left column:* Continuous normalized mean firing rate (FR) trajectories for isolated small, medium, and large motor unit populations. To illustrate precise temporal dynamics without population smoothing artifacts, these continuous traces and their associated trial-by-trial SEM envelopes are derived from a single representative participant. *Right column:* Population-level statistical distributions (*N* = 23) of the normalized mean firing rates extracted during the 500 ms execution epoch. Raincloud plots confirm that the pronounced spatial bias observed in the representative subject generalizes across the entire cohort, with highly significant, population-wide directional tuning preserved across all motor unit thresholds (Wilcoxon signed-rank tests; *** *p* < 0.001 across all unit sizes).

We hypothesized that this apparent peripheral silence might be an artifact of the gross EMG interference pattern, which heavily biases toward larger, execution-related motor units. To resolve this, the independent spike trains of putative small, medium, and large-amplitude motor units. This amplitude-based thresholding unmasked a striking functional divergence within the motor pool. Small, low-threshold motor units exhibited a selective, sustained increase in firing rate throughout the delay period for In-RF trials—ramping well before go cue and movement onset—while remaining entirely quiescent during Out-RF trials (target at +4.5°, −11°) (**Figure 3B**). In contrast, medium and large motor units showed virtually no preparatory delay activity.

Having established the presence of spatially tuned preparatory drive in low-threshold units, we next investigated the execution phase of movement to quantify distinct spatial tuning at the level of gross muscle output (0.00 to 0.50 s post-movement onset). Across the cohort (*n* = 23), evaluation of raw surface electromyography revealed substantial absolute amplitude differences based on movement direction (In-RF: Median = 10.70, IQR = 16.95 vs. Out-RF: Median = 3.99, IQR = 6.69 raw units). To rigorously account for inter-subject impedance variability, these signals were normalized to peak task activation. Consistent with the raw traces, normalized gross RMS EMG exhibited a pronounced, highly significant population-wide directional amplification (In-RF: Median = 0.68, IQR = 0.16 vs. Out-RF: Median = 0.29, IQR = 0.29 normalized units; Shapiro-Wilk test, *p* < 0.001; Wilcoxon signed-rank test, *W* = 269.00, *p* < 0.001).

To determine whether this robust peripheral spatial bias is driven by a uniform or size-dependent central recruitment strategy, we evaluated the isolated neural populations during the same 500 ms execution epoch. To ground our analysis in absolute physiological terms, we first evaluated the raw discharge characteristics of the motor pool. Raw firing rates scaled predictably across the size hierarchy and exhibited clear spatial preferences. During active In-RF movements, population-level raw firing rates reached a median of 35.33 spikes/s (IQR = 15.13) for small units, 19.73 spikes/s (IQR = 24.16) for medium units, and 12.64 spikes/s (IQR = 19.83) for large units. Conversely, Out-RF movements elicited substantially lower raw discharge rates across all populations (Small: Median = 12.96, IQR = 18.14 spikes/s; Medium: Median = 4.33, IQR = 13.79 spikes/s; Large: Median = 1.71, IQR = 10.24 spikes/s). Because absolute firing rates exhibit natural inter-subject variance and size-dependent scaling, continuous firing rates were subsequently normalized to each unit’s peak execution frequency. Within this standardized framework, all three motor unit populations exhibited powerful, spatially selective discharge rates. Small motor units displayed highly robust statistical tuning, with normalized firing rates nearly tripling during preferred movements (In-RF: Median = 0.94, IQR = 0.40 vs. Out-RF: Median = 0.35, IQR = 0.48 normalized units; Wilcoxon signed-rank test, *W* = 272.00, *p* < 0.001).

This strong directional bias was preserved across higher-threshold populations. Medium motor units exhibited a highly significant tuning profile (In-RF: Median = 0.79, IQR = 0.96 vs. Out-RF: Median = 0.17, IQR = 0.55 normalized units; Wilcoxon signed-rank test, *W* = 272.00, *p* < 0.001). Similarly, large motor units maintained a clear spatial preference, discharging preferentially for In-RF trajectories (In-RF: Median = 0.67, IQR = 1.05 vs. Out-RF: Median = 0.09, IQR = 0.54 normalized units; Wilcoxon signed-rank test, *W* = 267.00, *p* < 0.001).

### Preparatory modulation predicts execution dynamics exclusively within low-threshold motor unit networks

Having established that preparatory drive selectively targets low-threshold motor units, we next asked whether changes in this preparatory state actively dictate subsequent execution dynamics. Evaluation of raw discharge characteristics confirmed that small motor units uniquely exhibit sustained, ramping activity during the preparatory delay period—well in advance of the Go cue and overt movement. In a representative participant, this pre-movement drive was clearly quantifiable in absolute firing rates, which were robustly amplified following active neuromodulation (Pre-stimulation: Median = 3.96, IQR = 6.16 spikes/s vs. Post-stimulation: Median = 5.11, IQR = 4.64 spikes/s). Upon movement onset, this activity rapidly accelerated into vigorous execution-phase discharge (Pre-stimulation: Median = 52.82, IQR = 4.26 spikes/s vs. Post-stimulation: Median = 54.19, IQR = 2.42 spikes/s). To rigorously account for baseline inter-subject variance and enable unbiased comparisons of these recruitment dynamics across the motor pool size hierarchy, all continuous firing rates were subsequently normalized to each unit’s peak execution frequency. Furthermore, because single-unit trial distributions significantly deviated from normality (Shapiro-Wilk test, *p* < 0.001), exact non-parametric statistics were utilized for all within-subject comparisons.

Single-subject EMG recordings revealed a striking, size-dependent functional dissociation (**Figure 4A, B**). In a representative participant, active stimulation yielded a robust amplification of normalized preparatory firing rates in small motor units (Pre: Median = 0.069, IQR = 0.107 vs. Post: Median = 0.088, IQR = 0.080 normalized units; Wilcoxon rank-sum test, *W* = 386.00, *p* = 0.043). Importantly, this elevated preparatory baseline transitioned into an amplified execution state, where Post firing rates remained significantly higher than Pre (Pre: Median = 0.914, IQR = 0.074 vs. Post: Median = 0.938, IQR = 0.042 normalized units; Wilcoxon rank-sum test, *W* = 386.00, *p* = 0.043).

**Figure 4.**
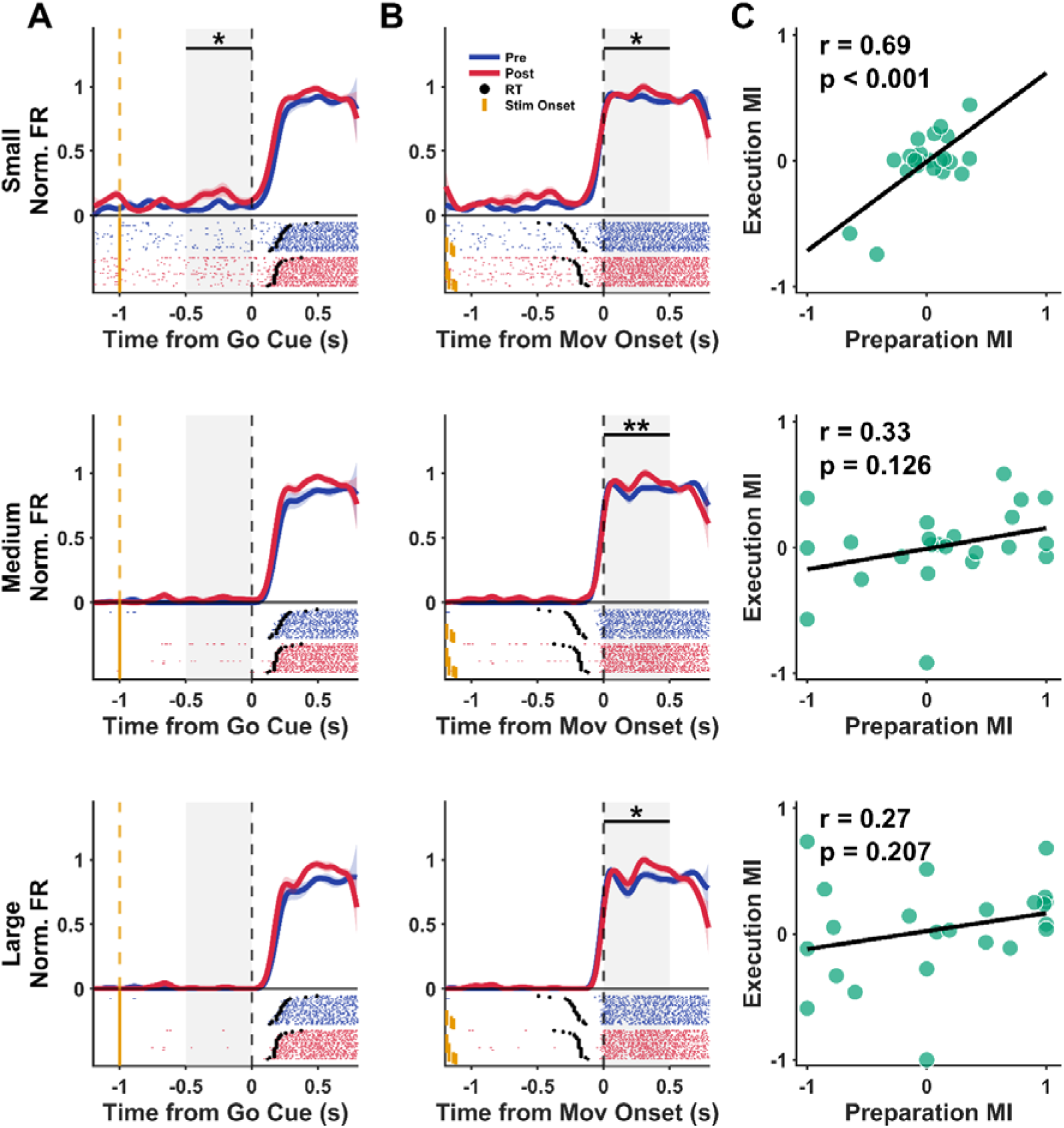
Preparatory neural modulation dictates execution dynamics exclusively within low-threshold motor unit networks. **(A)** Preparatory phase dynamics extracted from a representative participant. Continuous traces display the normalized mean instantaneous firing rate alongside trial-by-trial standard error of the mean (SEM) envelopes for isolated small (top), medium (middle), and large (bottom) motor units. Data are aligned to the Go cue (dashed black line at 0 s). The orange dashed line and corresponding raster ticks indicate the onset of onset of stimulus (−1.0 s), while black dots in the raster denote single-trial reaction times (RT). The shaded gray region delineates the 500 ms preparatory epoch (−0.50 to 0.00 s) utilized for statistical quantification. Significant tDCS-induced preparatory facilitation is observed exclusively in small motor units (* *p* < 0.05), whereas higher-threshold units remain unaffected. **(B)** Execution phase dynamics for the same representative units, realigned to movement onset (dashed black line at 0 s). The shaded gray region delineates the 500 ms active execution epoch (0.00 to 0.50 s). During overt movement, active stimulation significantly amplifies firing rates across the entire size hierarchy, encompassing small (* *p* < 0.05), medium (** *p* < 0.01), and large (* *p* < 0.05) motor unit populations. **(C)** Population-level predictive relationships (*n* = 23) between preparatory and execution states. Scatter plots correlate the Preparatory Modulation Index with the Execution Modulation Index, where positive values indicate a stimulation-induced increase in firing rate. A highly robust, predictive linear relationship is present exclusively within the small motor unit pool (Pearson correlation; *r* = 0.69, *p* < 0.001). This structural linkage is completely uncoupled in higher-threshold networks, with neither medium (*r* = 0.33, *p* = 0.126) nor large (*r* = 0.27, *p* = 0.207) motor units exhibiting significant preparatory-to-execution translation.

In stark contrast, higher-threshold units within the same participant were entirely decoupled from preparatory modulation. Both medium and large motor units remained functionally quiescent during the preparation window (Medium: Pre = 0.000 vs. Post = 0.000 normalized units, *p* = 0.096; Large: Pre = 0.000 vs. Post = 0.000 normalized units, *p* = 0.096). However, during active execution, these larger units exhibited highly significant, phase-locked amplifications (Medium Exec: Pre: Median = 0.873, IQR = 0.093 vs. Post: Median = 0.932, IQR = 0.075 normalized units; Wilcoxon rank-sum test, *W* = 355.00, *p* = 0.006. Large Exec: Pre: Median = 0.836, IQR = 0.126 vs. Post: Median = 0.916, IQR = 0.103 normalized units; Wilcoxon rank-sum test, *W* = 378.00, *p* = 0.027).

To determine if this size-dependent linkage represents a generalized principle of human motor control, we calculated a Modulation Index (MI) for both phases and assessed their linear relationship across the entire population (*n* = 23). Our analysis confirmed a highly specific, low-threshold channel for motor priming. In small motor units, preparatory MI strongly predicted execution MI (Pearson’s *r* = 0.687, *p* < 0.001). This predictive relationship was entirely abolished in higher-threshold pools, with neither medium motor units (*r* = 0.328, *p* = 0.126) nor large motor units (*r* = 0.273, *p* = 0.207) exhibiting a significant linear correlation.

## Discussion

To the best of our knowledge this is the first attempt at investigating the peripheral signature of motor preparation induced by neuromodulation of the supplementary motor area (SMA). In a randomised delayed reaching task, anodal stimulation of the SMA enhanced both motor preparation and downstream execution, shortening reaction times. These findings demonstrate that preparatory activity is also enhanced in the motor unit activity, whereas the transition into the planning and execution phase, particularly during movements with greater peak velocity, recruits and modulates medium- and higher-threshold motor units (medium and larger) more strongly. This progressive recruitment pattern suggests that SMA-targeted anodal-tDCS augments early preparatory neural drive while simultaneously facilitating the higher-force activation required for improved movement execution. Collectively, the converging behavioural and EMG evidence reveals that SMA neuromodulation amplifies a peripheral “preparatory set” (Carlsen et al.,2015).

### Relation to previous work

Prior work has established that neurons in the SMA exhibit delay-period activity that anticipates movement and distinguishes preparation from execution, since the neurons show little or no activity in the latter phase. These studies have framed the SMA as a controller of the preparatory set that biases motor circuits toward imminent action (Tanji, 1984; Tanji, 1994). In addition, prior EMG studies have mostly concluded that preparation was centrally confined because no overt muscle activity was detected during delays, a conclusion shaped by the sensitivity limits of single-channel EMG, coarse filters, and a focus on gross agonist activation rather than low-amplitude unit behaviour (Tanji & Evarts, 1976) that was hard to distinguish from baseline EMG noise. Subsequent behavioural neuromodulation work showed that increasing SMA excitability with anodal tDCS can shorten reaction times in a polarity-dependent manner, consistent with shifting the preparatory state closer to an initiation threshold (Carlsen et al., 2015 and others). Together, these strands suggested a cortical preparatory mechanism with behavioural relevance but lacked a clear peripheral signature at the motor unit scale, which we have tested in this study.

One potential limitation in previous EMG related studies which tested muscle activity during motor preparation, was the use of conventional EMG involving electrodes spaced at long distances which would have filtered small, local voltage fluctuations caused by activation of smaller motor units (Tanji and Evarts,1976). In addition, the use of standard signal processing approaches that emphasizes the analogue force related EMG activity such as RMS causes temporal low pass filtering of the EMG signal, emphasizing the whole muscle activity over earlier and smaller preparatory phase of the EMG signal. Furthermore, the use of more distal musculature involving wrist muscles ((Tanji and Evarts, 1976; Tanji & Kurata, 1982,1985), as well as using tasks involving cursor/joystick motion might have restricted the extent of physical movement as well (Prut and Fetz,1999), and produced delayed activity relative to reach onset due to the well-known temporal gradient of proximal to distal muscle related neural activity seen in the motor cortex (Jeannerod, 1981).

These potential limitations have been addressed in the recent work using high-density surface EMG (Rungta and Murthy, 2023). They reported context-specific early recruitment of small amplitude motor units during the delay, topographically tuned to target direction and predictive of reaction times, thereby revealing a peripheral correlate of central preparation that remains subthreshold for overt movement. Methodologically, their use of dense grids, bipolar derivations, and unit-scale spike detection overcame limitations of earlier EMG approaches that averaged away sparse, low-threshold firing which can also be observed in the current study by comparing Figure 4A and B. Conceptually, they argued that a modest, spatially specific increase in descending drive can selectively bias small motor units, preconfiguring the neuromuscular apparatus for rapid initiation without violating no movement constraints during the hold period. These conclusions converge with the findings of (Mellah et al.,1990), who first showed the activation of motor units that are selective during motor preparation but not during the execution of delayed movements in monkeys using needle EMG electrodes.

### Mechanisms underlying stimulation induced changes in the motor periphery during the delayed period

The current findings converge with and extend (Rungta and Murthy, 2023) on three fronts. First, behaviourally, anodal SMA stimulation further shortened reaction times in prepared/delayed trials (hold time), aligning with a causal role for SMA in configuring a launch-ready state. Second, preparatory firing rate increased (see figure 4A) RF-specific small motor unit discharge rose during the delay (see figure 3B), closely mirroring the early recruitment signature described by (Rungta and Murthy) but now under targeted SMA neuromodulation. Third, during execution, medium/large units showed the most prominent gains, consistent with recruitment principles as described by Henneman (1957) and Henneman et al. (1965), whereby enhanced central drive propagates to higher threshold pools once initiation occurs (see figure 4B ). This pattern links SMA level causal manipulation to the specific peripheral phenotype predicted by Rungta and Murthy, directly bridging central preparatory dynamics and neuromuscular readiness.

SMA stimulation in this study produced three phase-specific effects: selective RF- specific elevation of delay period activity particularly seen in small -motor unit activity (see figure 3B) with unchanged baseline, faster initiation, and disproportionately greater execution phase recruitment of medium/large motor units. Within Rungta and Murthy’s framework, the delay period increase, particularly in small unit activity indexes a peripheral preparatory set as these motor units are- context-specific that encodes future kinematic or postural states of the muscle and distinguish In-RF vs. Out-RF (see figure 3B), and a bias that predicts reaction time, thereby explaining the rise in preparatory without movement and a shift toward higher threshold- recruitment during movement execution (see Figure 4A,B). Extending this model to circuitry, the findings are consistent with the SMA and premotor cortex being a critical part of the preparatory subspace which, being movement null, prevents activation of the primary motor cortex (Kauffmann et al., 2014). This notwithstanding, we suggest that preparatory neural subspaces may encode desired future postures of relevant muscles (Kaufmann et al., 2014), which are conveyed possibly via reticulospinal pathways to spinal gamma motor neurons (Kearsley et al.,2021), yielding early, low-threshold motor-unit activity during the preparatory phase (Rungta & Murthy, 2023; Mellah et al.,1990). Although such gamma activation on its own would not cause muscle contraction, it could underlie a preparatory set in muscles that subsequently modulates alpha motoneurons via spindle-based feedback, consistent with equilibrium-point/referent-configuration models (Feldman, 1966; Feldman, 1986; Latash, 2010; Bizzi et al., 1982) but limited to the initial posture. This latter feedback could be the basis of both shorter reaction times as well as faster muscle contraction, leading to increased vigour of movement we observed (Shadmehr et al., 2019). (Hasnain et al.,2025) in a recent study support this division by showing that preparatory and execution components reside in separable neural manifolds that couple to distinct descending channels, offering a principled bridge from cortical subspace dynamics to the peripheral signatures observed here. This layered pathway explains the pattern observed here: delay period- small motor unit engagement, faster initiation, and amplified execution phase- recruitment, and predicts that selective modulation of reticulospinal drive will scale preparatory EMG topographies and movement vigor, whereas corticospinal activation will directly increase alpha motor unit activity via -high threshold motor unit recruitment leading to muscle contraction.

### Methodological Issues

The apparent absence of preparatory muscle activity in the previous work can be reconciled with the present and (Rungta and Murthy, 2023) results by considering sensitivity, resolution, and the signal processing analysis that was used. EMG from large and well separated electrodes with coarse temporal smoothing is ill-suited to detect sparse, low amplitude discharges of small units during movement preparation; averaging across channels and trials further suppresses these signals. In contrast, high-density grids (HD-sEMG), bipolar spatial filtering of electrodes that are 5mm apart, spike scale detection, and windowed analyses using peristimulus time histograms, centred on the delay capture subtle, context-specific changes that do not manifest as bulk activation. Thus, the older inference that “preparation is purely central” largely reflects measurement constraints rather than a true absence of peripheral modulation. Nevertheless, the current study is not without its limitations. The electric field targeting SMA was not individualized and is the spread to adjacent medial premotor regions cannot be excluded. Activation of the anterior pre-SMA, which is an inhibitory node, may in fact had the unintended consequence of suppressing muscle activation. The motor unit class attribution relied on a threshold amplitude rather than full decomposition and longitudinal unit tracking was not attempted. As such, these threshold crossing are still putative motor units and motor unit size maybe conflated with distance of the motor units from the electrode grid. Further, potential confounds such as expectancy from having a fixed hold time, and subtle posture changes during the hold time were controlled indirectly but not exhaustively. However, despite these potential limitations our results and conclusions and interpretations are still valid.

## Summary and Conclusions

While cortical delay period activity in SMA neurons defined the origin of the preparatory set (Tanji and Evarts,1976), polarity-dependent RT modulation established a causal leverage at SMA (Carlsen et al.,2015). The measurement of motor unit excitability indicated that preparatory drive can influence peripheral elements without overt movement, localizing this influence to selective, context-specific recruitment of smaller motor units detectible with high-density surface EMG (Mellah et al.,1990, Rungta and Murthy.,2023). The present study unifies these strands by showing that targeted SMA neuromodulation enhances precisely this peripheral preparatory signature and accelerates initiation, providing a mechanistic bridge from cortical preparation to neuromuscular readiness and its distinct relationship to subsequent execution.

## Author Contributions

**Ashutosh Kumar:** Conceptualization, Methodology, Investigation, Formal Analysis, Data Curation, Visualization, Writing – Original Draft, Writing – Review & Editing.

**Ayusha Desai:** Investigation, Writing – Review & Editing.

**Kalpajyti Hazarika**: Software, Methodology, Writing – Review & Editing.

**Aditya Murthy:** Conceptualization, Methodology, Supervision, Project Administration, Writing – Review & Editing.

## Data and code availability

Data and analysis code will be made available upon reasonable request to the corresponding author, in accordance with institutional and ethical requirements.

## Ethics approval

Human research was approved by the Institutional Human Ethics Committee, Indian Institute of Science, Bengaluru **(No - 01/23.02.2023)**

## Acknowledgments

We thank all study volunteers for their time and patience during the experiments. We are grateful to Ankita Das for her help with subject preparation and data collection. This work was supported by a grant from the Science and Engineering Research Board/ Anusandhan National Research Foundation (SERB/ANRF), Government of India, and additional intramural funding from the Indian Institute of Science (IISc).

